# Migration dynamics of an important rice pest: the brown planthopper (*Nilaparvata lugens*) across Asia – insights from population genomics

**DOI:** 10.1101/2020.03.07.981894

**Authors:** J. P. Hereward, X.H. Cai, A. M. Matias, G. H. Walter, C. X. Xu, Y. Wang

## Abstract

Brown planthoppers (*Nilaparvata lugens*) are the most serious insect pests of rice, one of the world’s most important staple crops. They reproduce year-round in the tropical parts of their distribution, but cannot overwinter in the temperate areas where they occur, and invade seasonally from elsewhere. Decades of research has not revealed their source unambiguously. We therefore sequenced the genomes of brown planthopper populations from across temperate and tropical parts of their distribution and show that the Indochinese peninsula is the major source of migration into temperate China. The Philippines, once considered a key source, is not significant, with little evidence for their migration into China. We find support for immigration from the west of China contributing to these regional dynamics. The lack of connectivity between the Philippines and mainland China explains the different evolution of Imidacloprid resistance in these populations. This study highlights the promise of whole genome sequence data to understand migration when gene flow is high – a situation that has been difficult to resolve using traditional genetic markers.

## Introduction

Understanding the migration dynamics and connectivity of populations is crucial to setting management priorities for pest insects. This is due to the central role population connectivity plays in the evolution of traits such as insecticide resistance and biological traits that enhance the pest status of an insect, such as the ability to overcome crops bred to be resistant to insect damage (Hendry *et al*. 2011). Migration is often the pathway by which insecticide resistance spreads (Denholm *et al*. 2002, Pasteur & Raymond 1996, Raymond *et al*. 1991). Genetics provides a way to assess migration dynamics in pest insects that cannot be tracked over large distances, but this has been difficult in the past when rates of geneflow (migration) between populations are high. The use of only a few genetic markers has often failed to provide the necessary resolution in high gene flow systems (Reitzel *et al*. 2013). Perhaps the ability to generate whole-genome sequence data can provide the necessary resolution in these high gene flow pest systems.

Rice is one of the world’s central food crops and is essential to global food security (Long-ping 2014). One of its most serious threats, the brown planthopper (*Nilaparvata lugens*) (Stål) (Hemiptera: Delphacidae) (Matteson, 2000, Sogawa & Cheng, 1979), physically damages rice plants with its phloem feeding and also causes indirect damage through virus transmission (Nault & Ammar, 1989). Despite an extraordinary amount of research, the migratory routes and population connectivity of this pest remain poorly understood (Bao *et al*., 2000, Cheng *et al*., 1979, Dung, 1981, Jiang *et al*. 1982, Kisimoto, 1976, Kisimoto & Sogawa, 1995, Riley *et al*., 1991, Rosenberg & Magor, 1983, Seino *et al*., 1987, Sogawa & Cheng, 1979). A clear understanding of these aspects of migration is essential to managing this important pest species effectively, and has been the core focus of decades of research. The general consensus is that they travel from the tropical part of their distribution into the northern regions of Vietnam and the southern, central and eastern parts of China, migration from these areas follows, apparently associated with the spring monsoon into south Korea and Japan; and a reverse migration to the tropics, with the autumn monsoon, has been proposed (Cheng *et al*., 1979, GAABPG, 1979, Jiang *et al*. 1982, Kisimoto, 1976). The primary aim of this study was to address this important question using whole-genome population genetics and to determine the connectivity of potential southern sources and Chinese populations of brown planthoppers.

Brown planthoppers are present throughout the year in the tropical parts of their geographical distribution, which includes the Indochinese peninsula, Southeast Asian islands and parts of India (Fig. 1). Here, conditions are ideal for *N. lugens*, where they generally complete one generation every 30 days (Li, 1984, Zhu *et al*., 2000), resulting in 12 to 13 generations per year. These bugs cannot, however, overwinter on rice in the northern parts of their distribution (where the coldest monthly temperature is below 12 °C), so seasonal appearances in their northern range is a result of migration (Cheng *et al*., 1979). Migration of brown planthoppers commonly starts in March, culminating in a peak immigration into the Yangtze river basin in June to July, where the insects may persist for up to five generations (Cheng *et al*., 1979, Ding, 1979, Hu *et al*., 2014).

**Fig. 1.**
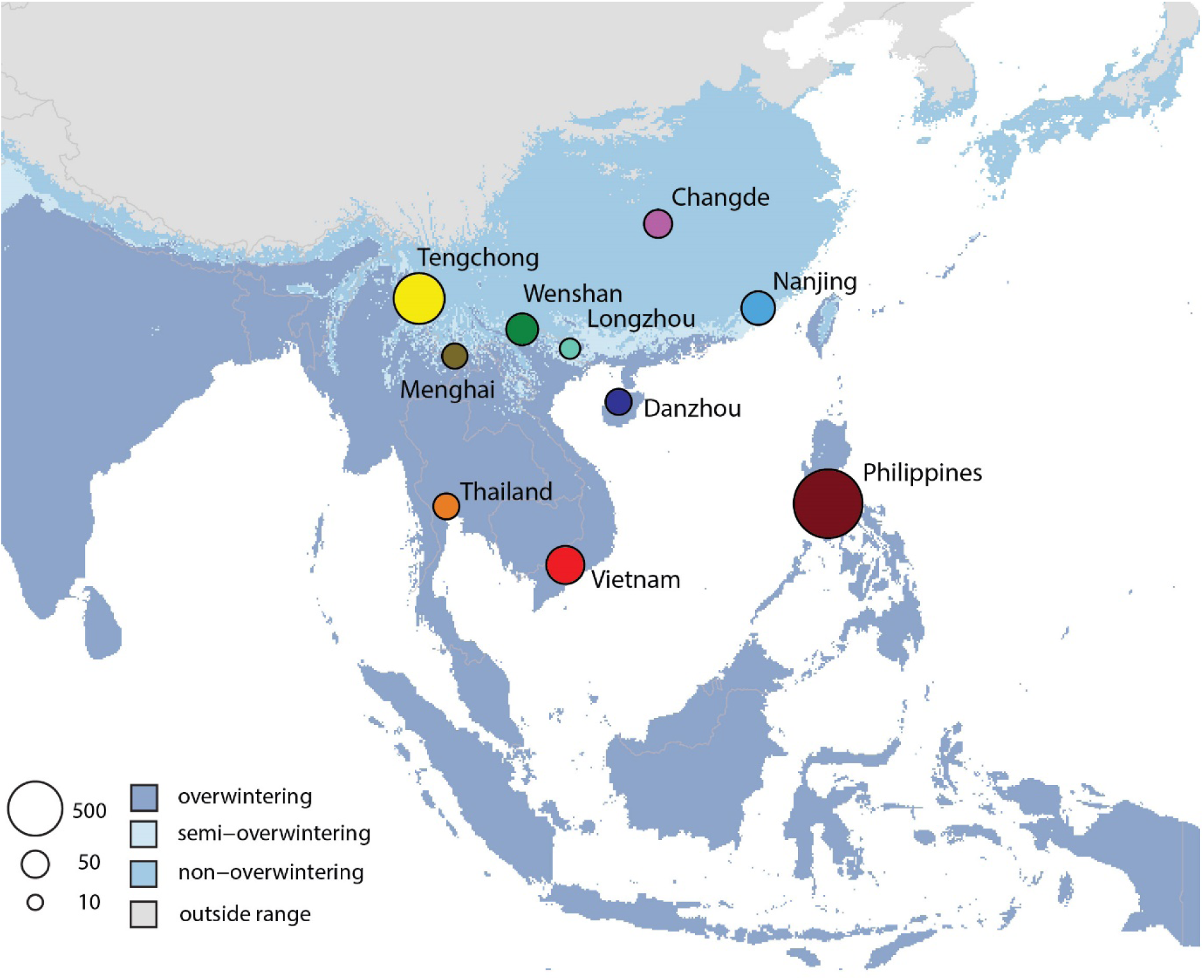
Map showing the geographical range of brown planthoppers and the extent of the overwintering and non-overwintering parts of its range (see key, bottom left). Populations sampled in this study are shown and the size of the circles represents the number of population specific (private) alleles in each population.

Numerous studies have investigated seasonal long-distance movement of this species through an extensive assortment of techniques: catching individuals at sea, in the air with airplanes, and on mountaintops (Cheng *et al*., 1979, Dung, 1981, Kisimoto, 1976, NCRGBP, 1981). Meteorological analyses, radar observations, and mark-recapture experiments have also been conducted (Riley *et al*., 1991, Rosenberg & Magor, 1983, Seino *et al*., 1987). Based on these studies, there is a widely held belief that migration occurs from southern parts of their distribution to the Chinese mainland, south Korea, and Japan (Bao *et al*., 2000, Kisimoto & Sogawa, 1995, Sogawa & Cheng, 1979).

Three putative sources of planthopper migration into China, south Korea and Japan, have been identified and these are not seen as mutually exclusive. The Indochinese peninsula is generally considered the primary source (Otuka *et al*., 2008, Wu et al., 1997, Zhu *et al*., 1994). In this region, the Red River and Mekong River Deltas are planted extensively to rice. The Mekong Delta alone contains roughly 10,000 square kilometres of rice, and supports up to three crops annually (Nguyen, 2007). The smaller Red River Delta is limited to two crops per year, presumably supporting fewer brown planthoppers. A putative migration route has origins in the Mekong River Delta, moving to the Red River Delta in February and March, and then to south China, east China and Korea and Japan in the months that follow (Wu *et al*., 1997). However, independent analyses of the seasonal monsoon suggest that brown planthopper populations in the Mekong River Delta cannot migrate into the Red River Delta and then to China because the seasonal winds in February to April are not suitable (Hu *et al*., 2017, Zhai & Cheng, 2006).

The second putative source is the Philippines where rice is also the main food crop. Weather simulations indicate that brown planthoppers could migrate from the Philippines into Taiwan and Japan, but also to southern China, sometimes in association with typhoons (Otuka *et al*., 2005). The Philippines populations possibly mix with Indochinese populations in southern China before arriving in Japan (Otuka *et al*., 2005, Otuka *et al*., 2012), but this has not been tested further. Various features of the Philippine bugs suggest that they are different from the mainland populations. They are nearly one hundred times more susceptible to the pesticide imidacloprid than populations in Taiwan, northern Vietnam, China, and Japan (Matsumara *et al*., 2008). In addition, the Philippines and other east Asian populations of brown planthopper differ from one another in their ability to overcome planthopper-resistant rice varieties (Sogawa 1992).

The third putative source is Myanmar and Bangladesh, but these bugs are likely to extend only into the west of Yunnan province (China) and it is thought that they would not be able to migrate into central and eastern China because of mountainous barriers and unsuitable monsoon conditions (Wu *et al*., 1997). These three putative sources of brown planthopper migration have found varying levels of support from empirical studies, but have not been tested directly.

Studies based on mitochondrial DNA (mtDNA) indicated high gene flow across the distribution of brown planthopper (Mun *et al*., 1999). One haplotype was found in every country sampled across South and East Asia, and since only three haplotypes were recovered from the 71 individuals investigated little resolution of the migration dynamics was possible. Understanding population connectivity in this pest is key to understanding how undesirable traits like insecticide resistance will spread. The objective of this study was to investigate population connectivity of populations of brown planthopper at a broad geographic scale, throughout China and Southeast Asian, to better understand the migration routes of this key insect pest of rice.

## Materials and Methods

### Sample collection and sequencing

*Nilaparvata lugens* adults were collected in rice fields from 10 locations in China, Vietnam, Thailand and Philippines from April – August 2010 (Supplementary data table S1). The population from Los Baños, Philippines, was collected in 2009. Samples were collected directly into 95% ethanol and then stored at −20°C, prior to DNA extraction. In Tengchong, Nanjing (Fujian province) and Changde, brown planthoppers cannot overwinter at any life stage of development. In Menghai, Wenshan, and Longzhou they can sometimes be found in rice volunteers in winter. In the tropics (Vietnam, Thailand and Philippines) they survive throughout the year.

DNA was extracted from six individuals from each population using TaKaRa MiniBEST Universal Genomic DNA extraction Kit Ver.5.0 (U-Me Biotech, Wuhan). Library preparation and sequencing was conducted by Total Genomics Solution (Shenzhen, China). Sequencing was performed on Illumina HiSeq X Ten returning 150bp paired end reads with an insert size of approximately 280bp. Sequence data quality metrics were estimated with Fastqc (Andrews, 2018) and Fastqsceen (Winget, 2018). The sequence data of three of the individuals from Vietnam did not contain sufficient data that mapped to brown planthopper, after quality filtering, and were discarded from further analysis. Six additional individuals from that site (Vietnam) were sequenced with BGI genomics (Shenzhen, China) with the same insert size, read length, library preparation and sequencing technology. We discarded the three Vietnamese samples that failed quality control but kept the three from the original sequencing that had passed, resulting in nine individuals from this population. All other samples had high quality sequences and we used all six individuals in downstream analysis, resulting in full genome sequences of 63 individuals in total. We removed adaptors, and quality-trimmed the end of the sequences of all samples to q10 using bbduk from the bbtools package (version 36), with a kmer of 8 (Bushnell, 2018).

### Data assembly

We mapped all reads to the brown planthopper reference genome, assembly GCA_000757685.1 of bioproject PRJNA177647 in Genbank (NCBI), which is approximately 1.2 Gbp (Xue wt al., 2014), using BWA mem v0.7 (Li & Durbin, 2009). We followed GATK (Genome Analysis Tool Kit) best practices (Van der Auwera *et al*., 2013) and called variants using HaplotypeCaller (gatk v 4.0.10.1), this was made parallel by splitting the reference into windows, each containing complete contigs. For each window we called individual g.vcf files and later combined these and performed joint genotyping using GenotypeGVFs (gatk v 4.0.10.1).

We filtered *snp*s (single nucleotide polymorphisms) to quality Q30 using vcftools, then to a minimum depth of three (as studies have showed that with appropriate filtering genotypes can be called from as few as three sequence reads) and a maximum depth of 30 (three times our highest coverages). Following this we removed any marker that was missing more than 30% of the data (30% of individuals were not genotyped), to remove the markers most affected by missing data. We then removed any locus with a minor allele count below three (e.g. one homozygote and one heterozygote). Applying hard filters for minor allele frequency can cause weak structure to be missed (Linck & Battey 2019), but the vast majority of singletons are errors, and also confound analyses. We used the minor allele count (=3) instead, because it is the most conservative way of removing singletons, but not rare alleles. We repeat-masked the reference genome using RepeatMasker 4.0.8 using the Dfam and RepBase databases – the repeat-masked reference was converted to a bed file, and this was used to remove *snp*s in repetitive regions of the genome with vcftools (Danecek *et al*., 2011). We then created a bed file with all the contigs below 5kbp and removed all of these *snp*s in the same way, assuming that some of the smaller contigs were likely to be less reliable. All indels and non-biallelic *snp*s were removed leaving only biallelic *snp*s, due to the requirements of downstream analyses. We filtered out markers that were out of Hardy-Weinberg Equilibrium because most of these are likely errors, and we were interested in the neutral process of migration, rather than investigating any effects of selection. Deviations from HWE were calculated separately using vcftools in the tropical populations Philippines, and Indochinese Peninsular (Thailand and Vietnam combined, see below), and then all markers with deviations from HWE in these populations were removed from the total dataset (using a threshold of p=0.05). This final dataset contained 762,576 *snp*s and is referred to as the *snp* genotype data. An alternative approach to calling *snp*s and filtering them is to infer genotype likelihoods, with some analyses methods then being able to use the likelihoods rather than the raw genotypes themselves, preserving the uncertainty in the inferences. We also called genotype likelihoods with the program ANGSD (Korneliussen *et al*., 2014), using the Samtools likelihood model; this dataset is referred to as the genotype likelihood data.

### Mitogenome analysis

We mapped the sequence data to the mitochondrial genome of *N. lugens* using BWA (Burrows-Wheeler Aligner, Li & Durbin, 2009). There are six mitochondrial genomes for brown planthopper on Genbank, however only one has the control region and repeats (JX880069). This mitogenome also had a large insertion between *ND2* (NADH dehydrogenase two) and *COI* (cytochrome oxidase subunit one) that was not supported by read-mapping, so we removed this region (based on alignment to the other five mitogenomes) and used this edited sequence as the reference. We used a pipeline to call *snp*s and prepared a consensus sequence for each individual. Briefly, after mapping, we removed the unmapped reads using samtools (Li *et al*., 2009), sorted, marked duplicates and indexed the bam file using GATK4 (Van der Auwera *et al*., 2013), called *snp*s using HaplotypeCaller from GATK4 in ploidy 1 mode, filtered the vcf file to q30 with vcftools (Danecek *et al*., 2011), made a consensus sequence with bcftools (Li *et al*., 2009), masked regions with no mapping using bedtools (Quinlan & Hall, 2010), and then renamed the fasta file with the name of the sequenced individual in a perl wrapper script.

We manually checked the resulting consensus sequences by mapping back the raw reads, and had to manually edit a small portion of the *ND2* gene because read mapping in that region indicated some kind of duplication, so we edited based on the reads that did not disrupt the coding sequence of the gene. The mapping was also unreliable around the repetitive control region in the reference, so we deleted this from the overall alignment. Sequences were aligned with MAFFT (Katoh & Standley, 2013) and we made a TCS haplotype network with Popart (Leigh & Bryant, 2015).

### Genetic structure

The distribution of brown planthoppers was plotted on a map, areas with mean monthly temperature of the coldest month below 13.5°C were designated as non-overwintering, between 13.5°C and 15°C were designated as intermediate, and over 15°C as overwintering. The temperature data were obtained from WorldClim global climate layers (https://www.worldclim.org/) accessed using the R (R Core Team, 2017) package raster.

We first examined the overall population structure by calculating F_ST_ (Weir and Cockerham, 1984) using the full *snp* genotype data described above. Overall F_ST_ was calculated in hierfstat (Goudet, 2004), as were pairwise F_ST_’s, performing 1,000 bootstrap replicates. We used the genotype likelihoods to infer admixture proportions using NGS admix (Skotte *et al*., 2013), which is designed to infer admixture proportions in low coverage sequencing data.

A discriminant analysis of principal components (DAPC) (Jombart *et al*., 2010) was performed on the full *snp* genotype data to assess the genetic relationship between the populations. DAPC maximises differences between defined populations across multiple principle components. Cross-validation indicated that 30 components should be retained. The DAPC analysis was performed using the adegenet package in R (Jombart, 2008).

To assess population structure without pre-defining populations we performed a principal component analysis (PCA) on the full *snp* genotype data. Principal component analysis cannot be performed on data with missing values, so for each population we replaced missing data with the population specific mode using custom R scripts (see data availability).

### Assignment testing

We used assignment testing to infer the source of the immigrants into the non-overwintering locations. Based on the PCA and admixture analyses we designated the “core - overwintering” populations (including Tengchong), which we assume to be the putative sources, into three groups; Philippines, Tengchong and Indochinese peninsular. Indochinese peninsular comprised of Thailand and Vietnam. We generated linear discriminant functions based on these three groups, using DPAC (Jombart *et al*., 2010) and retaining 10 axes. We then assigned individuals from the non-overwintering sites to these three groups using DAPC (Jombart *et al*., 2010).

### Genetic diversity and population-specific alleles

The number of population-specific alleles in each population was calculated using a custom R script which calculated allele frequency and then identified *snp*s that had a value other than zero in only one population, these were represented as the size of the circle for each population on the map. We then tested whether the population-specific alleles were driving genetic structure by running PCA on the full *snp* data and on a dataset with all population-specific (private) alleles removed.

The genetic diversity summary statistics pi and theta were calculated from the genotype likelihood data for every 10 kb sliding window with 2.5 kb overlap across the genome in ANGSD (Korneliussen *et al*., 2014). This analysis aimed to capture the genomic variance – that is the variation in patterns across contigs or chromosomes. These data were subsampled by selecting one out of every three windows. Windows that were not available across all 10 populations were then discarded. Windows in which the effective number of sites was less than 100 bp were also removed. The remaining windows were then used to plot the genomic diversity. There was no variation across populations when considering the raw distributions. The estimate for each window was standardized by the mean and standard deviation across the population ((x-x^−^)/(sd(x))), with the direction (sign) of the numerator indicating higher or lower genomic diversity.

## Results

The mitogenome analysis indicated little spatial structuring of haplotypes. We recovered 53 unique haplotypes from the 63 different individuals, based on 296 variable sites in the 14,380bp mitogenome alignment. Of the few shared haplotypes, one was found in Wenshan, Menghai and the Philippines, and another in Menghai, Longzhou and Thailand (Fig. 2). When we restricted the mtDNA analysis to COI, the same major haplotype was present across all populations, but several unique haplotypes were found in the Philippines (Fig. 2).

**Fig. 2.**
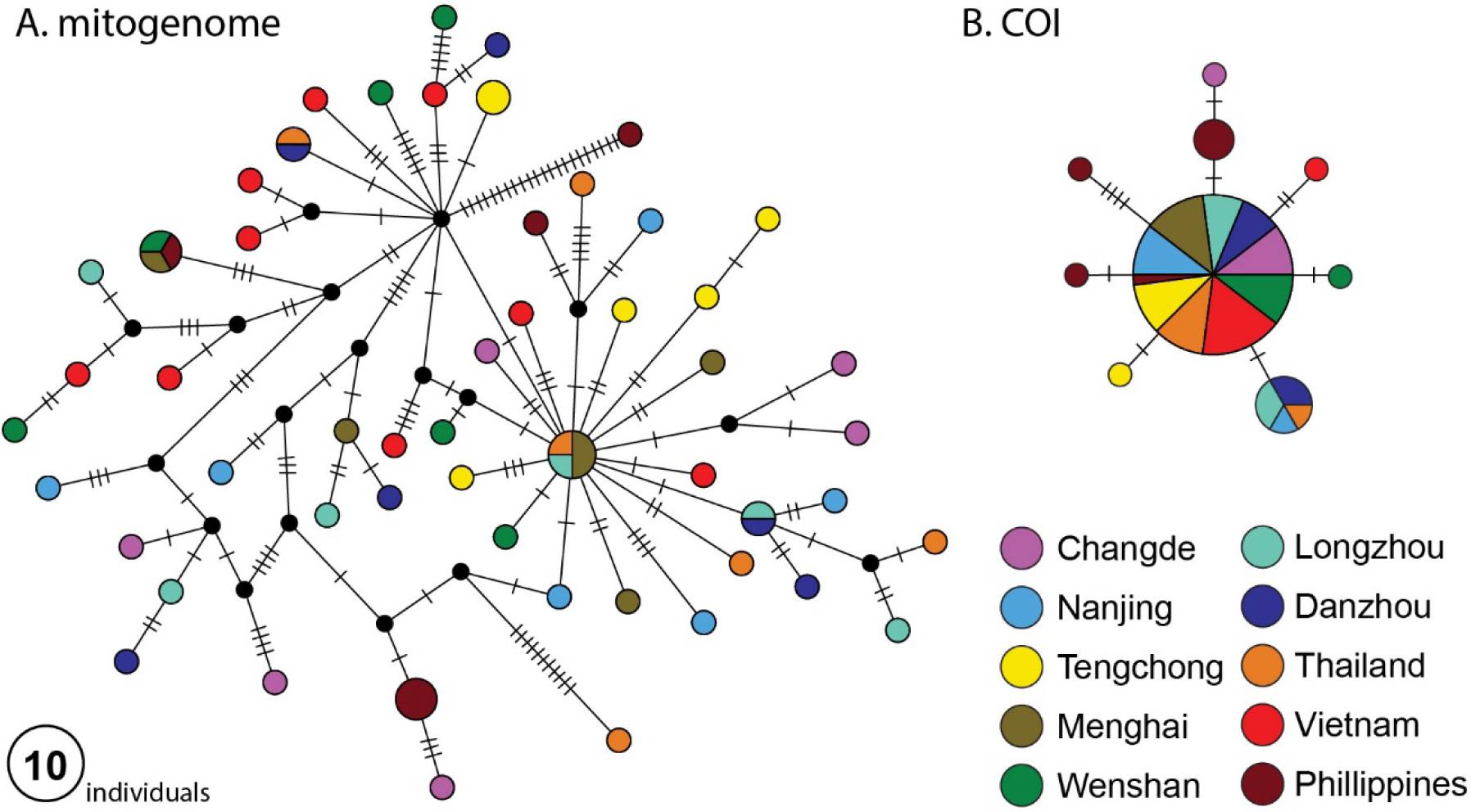
Mitochondrial genome analyses. Haplotype networks based on the complete mitochondrial genome (16,794 bp, **A**), and the COI gene only (2,795 bp, **B**) the size of each circle represents the number of individuals that have the same sequence. For localities see Fig. 1.

We achieved a mean coverage of 10.5x across the filtered *snps* and around 6x coverage across the whole genome in our initial mapping (Supplementary data table S2). Genetic differentiation was low across all populations, with an overall F_ST_ of 0.0093 (+-95% CI 0.0092 - 0.0094). The highest pairwise F_ST_’s were from comparisons of the Philippines to other populations (Supplementary table S3), and then from comparisons of Tengchong to other populations. Despite the low overall genetic differentiation, whole genome sequence data revealed major insights into the migration dynamics of brown planthopper. Admixture analysis (NGSadmix), DAPC, and the PCA indicated that the Philippines insects represent a distinct population with very little evidence of admixture from the Philippines into mainland populations (Fig. 3, Fig. 4). Tengchong, in the far west of China was also separated from other populations in the admixture analysis, with evidence of admixture across the rest of the mainland sites (Fig. 3). The PCA separated some individuals from the Tengchong population but others were placed in the main cluster, indicating a mixture between common genotypes and an unidentified source population.

**Fig. 3.**
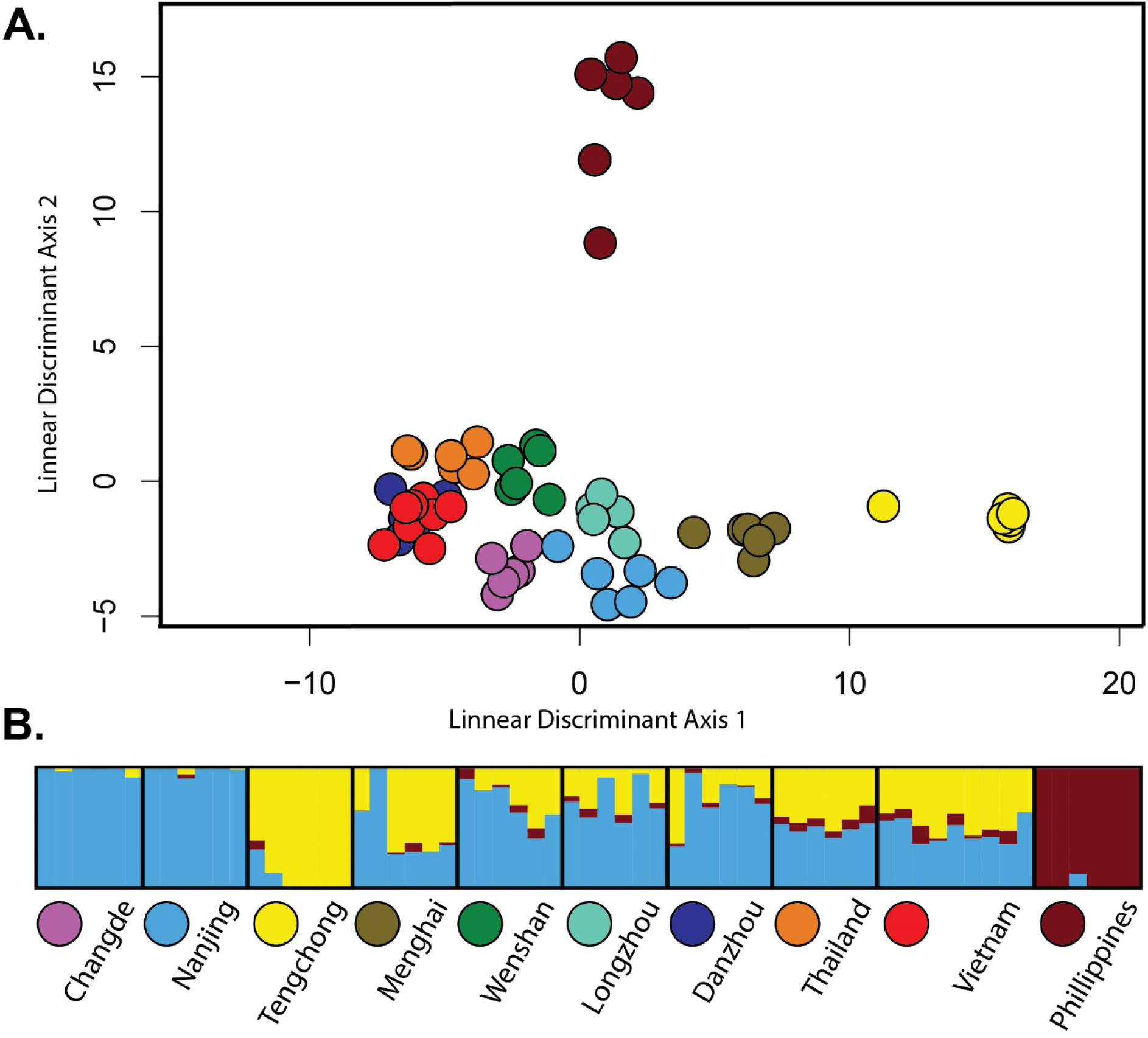
**(A.)** discriminant analysis of principal components (DAPC) of the *snp* genotype data maximising genetic differences across defined populations. **(B.)** Admixture analysis based on the genotype likelihood data using *NGSAdmix* to infer admixture proportions. Each bar is one individual and the colours represent the posterior probability of assignment to each of three putative ‘population’ clusters.

**Fig. 4.**
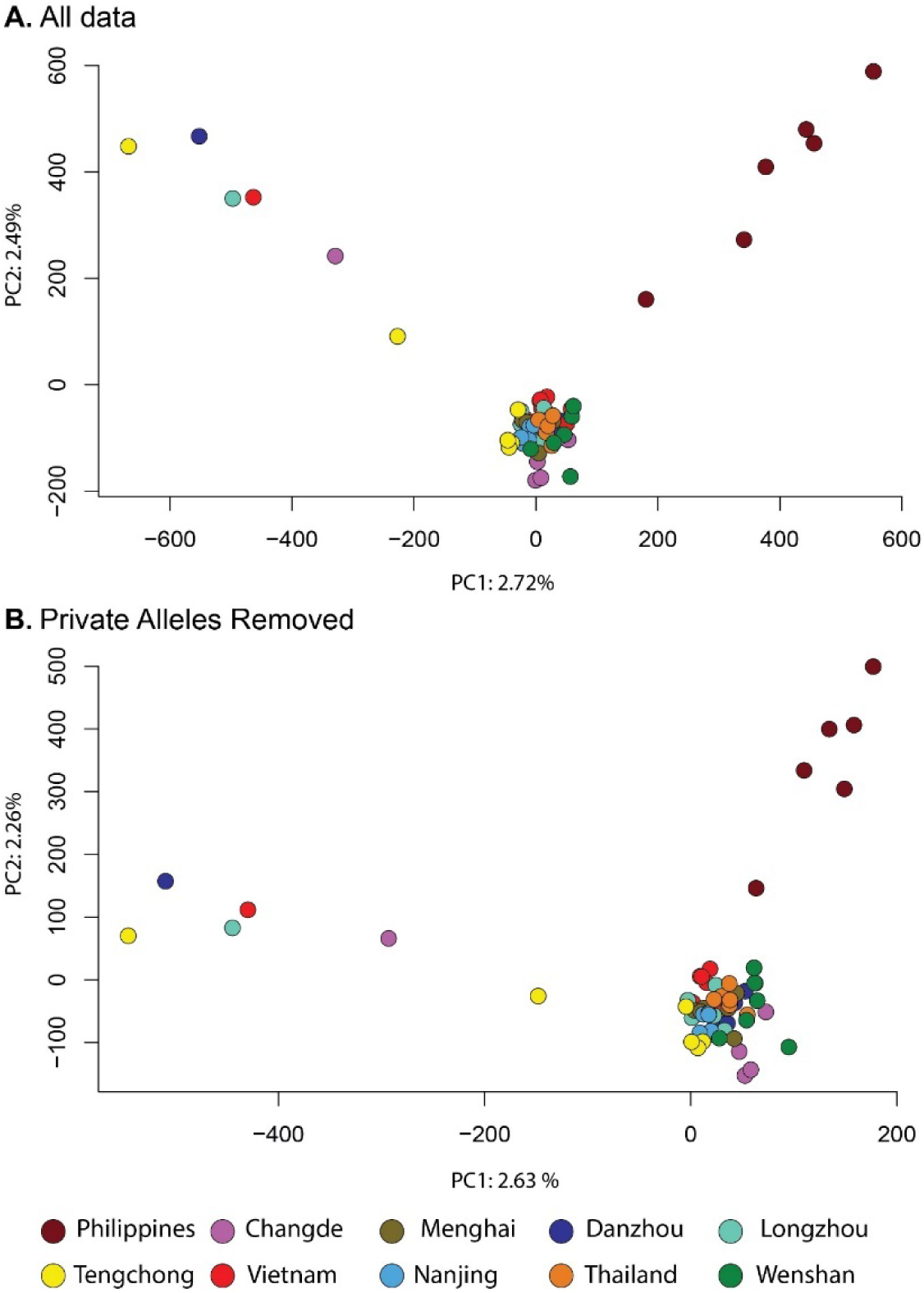
Effect of population-specific alleles on genetic structure. (**A**) Principal components analysis (PCA) of the full *snp* genotype data. (**B**) PCA of *snp* genotype data with all population-specific alleles removed. The genetic structure was not driven by private alleles.

The Philippines and Tengchong samples had more unique, population-specific alleles than the other populations (Fig. 1), even though the PCA indicated that Tengchong is a mixed population (Fig. 4). This likely indicates a distinct source of genetic diversity (presumably outside the geographic scope of this study) that contributes to the migration into Tengchong. We tested whether the population-specific alleles were driving our inferences of population structure by removing all of them and conducting a separate PCA analysis. The PCA remained qualitatively the same when all population-specific alleles had been removed from the dataset (Fig. 4, bottom), indicating that allele frequencies, rather than population-specific alleles, are driving our inference of genetic structure. The higher number of population-specific alleles in these two populations was not the result of higher overall genomic diversity as these two populations had lower genomic diversity than other populations (Fig. 5). The non-overwintering sites (Changde and Nanjing) did not have lower genomic diversity than the sites in tropical and subtropical regions that have year-round production of brown planthoppers (Fig. 5).

**Fig. 5.**
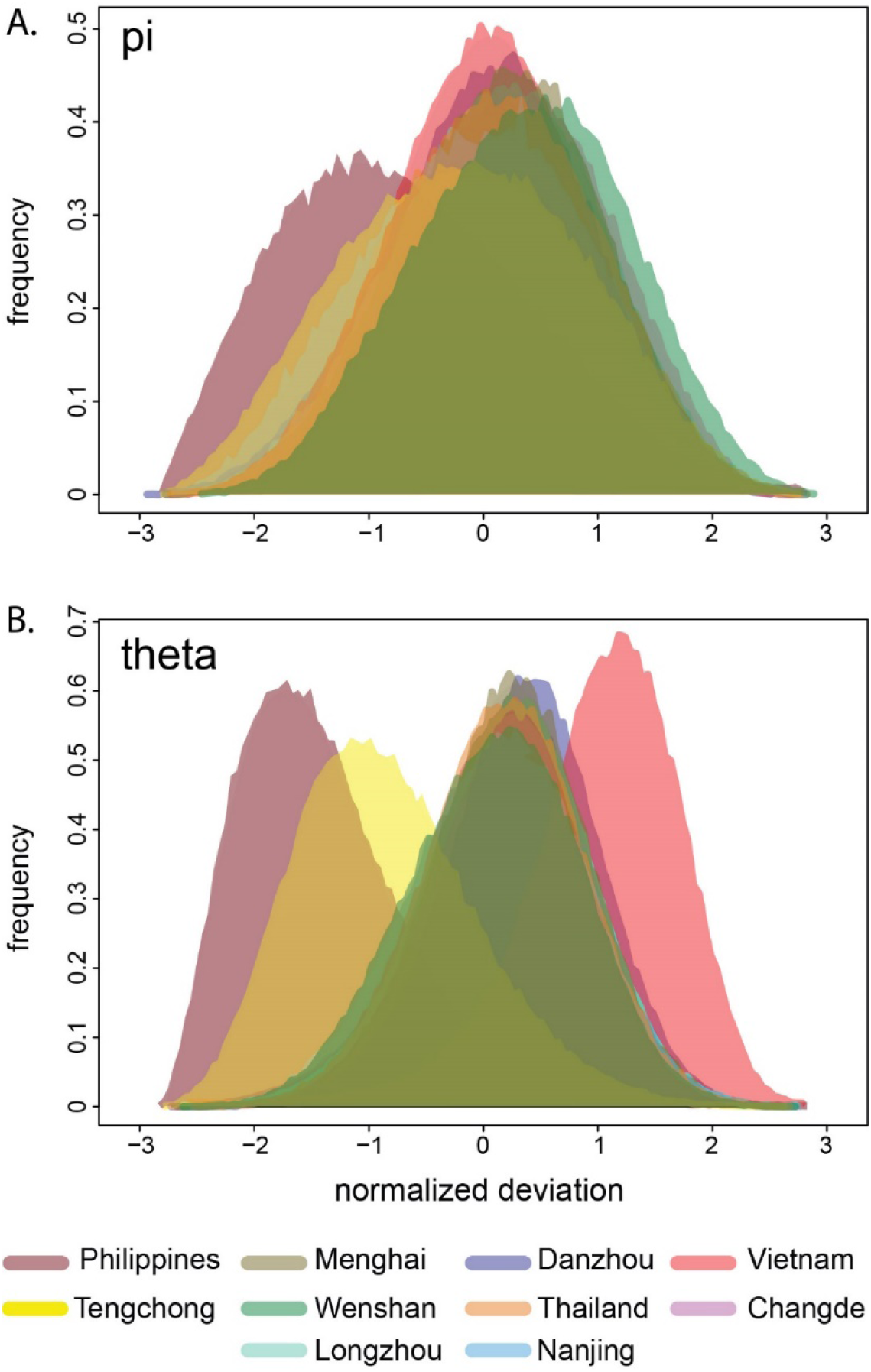
Genomic diversity. Plots of the normalised deviation from mean diversity measures pi (**A**), and theta (**B**), raw histograms shown.

All individuals from the non-overwintering sites were assigned to the Indochinese peninsular group (comprising Thailand and Vietnam) at close to 100% probability. When we plotted the linear discriminant scores for the non-overwintering individuals and the three “core-overwintering” groups, the non-overwintering individuals were placed between the Indochinese peninsular group and Tengchong (but closer to Indochinese peninsular). This is likely a result of the admixture between Tengchong and the other mainland populations (Supplementary Fig. 1).

## Discussion

Our results show that a population genomics approach contributes considerably to our understanding of the migration dynamics of brown planthoppers. The Indochinese peninsular is indicated as a major source of brown planthoppers in their migration to the temperate regions of China (and most likely onto Japan). The Philippines population was genetically distinct from the other populations in all analyses and little evidence of gene flow from the Philippines into temperate China was found. Tengchong (in western temperate China) also formed a separate group in the admixture analysis and DAPC, but the PCA indicated a mixed population (Figs. 3 & 4). The high prevalence of population-specific alleles in this population likely indicates immigration from more western regions (potentially Myanmar, Bangladesh and India, which were not sampled in this study). The populations in subtropical China and the tropical Indochinese peninsular show admixture with the Tengchong population, indicating that planthoppers from these putative western sources may contribute to the overall migration dynamics, despite the mountainous barriers and unsuitable monsoon conditions, and contrary to previous assertions (Wu *et al*., 1997).

Our mitogenome analysis is broadly in line with the results of Mun *et al*. (1999) who found only three haplotypes, one of which was present across all sites that they sampled in China, the Philippines, Bangladesh, Malaysia, Vietnam Thailand and Korea. By using the whole mitogenome we recovered many more haplotypes, but still no geographical clustering of the haplotypes (Fig. 2). This indicates no long-term separation of the various populations and is consistent with an interpretation that the insects are highly migratory and populations from all parts of its distribution have been connected through migration and gene flow in the relatively recent past. Leveraging genome-wide nuclear *snp* data, we were, however, able to make inferences into the more recent migratory dynamics of this species (as outlined above and below).

Physiological comparisons of bugs from temperate, subtropical and tropical regions have led to suggestions that tropical populations of brown planthopper do not contribute to the migratory dynamics of this pest (Iwanaga *et al*., 1987, Nagata & Masuda, 1980, Wada et al., 2007, Wada *et al*., 2009). Our analysis shows, by contrast, that the tropical populations in the Indochinese peninsular, the subtropical populations in China, and the temperate populations in China are connected genetically, all of which indicates that the tropical populations in the Indochinese peninsular (Vietnam and Thailand) are major sources of the migration into China. Planthoppers from these locations essentially form a single panmictic population, as revealed in the PCA, and they were separated only in the DAPC (which maximises differences across designated “populations”). We expected that the non-overwintering populations (Changde and Nanjing) might have reduced genetic diversity if they had experienced a founder effect following immigration in the spring. This was not the case (Fig. 5), and the same amount of genetic diversity was present in these two populations as in the Indochinese peninsular populations. This indicates that sufficiently large numbers of insects make the northern migration that bottlenecks or founder effects do not occur.

Population genetics approaches to estimating migration rates enable inferences to be made about small insects over the large distances that separate populations, and which are not possible with ecological approaches to measuring dispersal such as mark recapture techniques. Our study indicates that using the whole genome provides greater power to make these inferences than previous molecular methods based on relatively few markers (such as the mitochondrial analysis of Mun et al (1999)), as suggested by Luikart et al. (2003). It is not yet clear what the power difference is between empirical microsatellite datasets and whole genome analyses to detect migration because there are not many datasets to compare. Studies that compare microsatellites and *snps* generally find that moderate numbers of *snps* can detect the same genetic structure as microsatellite data, and that the resolution increases as the number of *snps* increase (Fischer et al 2017, Garke et al 2011). Generating whole genome sequence data also allows further analysis of endosymbionts and tests of selection in the future, although this was outside the scope of the current study.

This study highlights that it is still difficult to make precise estimates of migration using classic population genetic methods when gene flow is very high (high migration systems like brown planthopper). The number of individuals that could be included in this whole genome analysis was limited by cost. Increasing the number of individuals per population may have provided greater power in our assignment testing. The real promise of whole genome sequencing in this kind of study, however, likely lies in the potential to perform not only parentage analysis, but to leverage the information from linkage and recombination to infer deeper pedigree information and estimate migration across multiple generations. This approach would require many individuals to be sampled from each putative population and sequenced with sufficient depth, and is currently cost prohibitive, but will likely become feasible with further advances in sequencing technology.

### Implications for management

The Philippines has been reported as a major source of the northward migration of brown planthoppers into temperate regions of China and Japan, based on trap catches, backwards telemetry and modelling (Otuka *et al*., 2005, Otuka *et al*., 2012). In our analysis the Philippine population of brown planthoppers formed a highly distinct cluster in the admixture analysis (Fig. 1) and the PCA (Fig. 4). There is very little evidence of admixture of the Philippine population into the temperate or subtropical Chinese populations. This indicates very little gene flow and migration between the Philippine populations and mainland Chinese temperate populations. Our results explain the difference reported between the Philippines and other populations in their resistance to imidacloprid (Matsumara *et al*., 2008), as well as their relatively different responses to new rice varieties (Sogawa, 1992). Recently, a distinct strain of the planthopper endosymbiont, *Arsenophonus*, has been associated with imidacloprid resistance (Pang *et al*., 2018). All susceptible insects in that study had a distinct strain of *Arsenophonus* and when this strain was transplanted into resistant individuals, they became susceptible. This strain is found predominantly in the Philippines, although it was also detected at a lower frequency in the Chinese mainland province of Guangxi (Pang *et al*., 2018). By showing that the Philippines population is distinct, with little evidence for migration between the Philippines and the Chinese mainland, our data explain the distribution of the susceptible and resistance related *Arsenophonus* strains, although the presence of a susceptible-related endosymbiont in Guanxi (China) in that study is most likely the result of rare immigration of Philippines planthoppers into mainland China. This highlights the key role of migration in the evolution and spread of insecticide resistance (Denholm *et al*. 2002, Pasteur & Raymond 1996, Raymond *et al*. 1991).

Our results show that the Philippines is distinct from the mainland populations and there is very little migration between them. Insecticide resistance traits that evolve in the Philippines are less likely to spread to the mainland and *vice-versa*, as seems to be the case for imidacloprid (Pang *et al*. 2018). Populations in mainland southeast Asia and mainland China are highly connected by gene-flow, however, and insecticide resistance that evolves in Vietnam or Thailand is highly likely to spread to China, and could quite possibly spread back the other way. This creates a situation where insecticide resistance management has to occur across borders. Attempts to manage insecticide resistance in China that do not consider the populations and insecticide selection pressures south of the border would be futile.

Our data also indicates that there may be a somewhat distinct population to the west of Tengchong, perhaps in India. Future studies should investigate populations further to the west of the insect’s distribution, from Japan, and more sites in the southeast Asian part of its distribution to further establish the migratory dynamics of this insect and the routes by which insecticide resistance traits travel.

## Conclusions

In summary, we used whole genomes to make new inferences about brown planthopper migration. Chinese and Indochinese peninsular populations should be considered the same in terms of management and insecticide resistance. The Philippines population is distinct from these mainland populations and this explains the prevalence of imidacloprid susceptibility there compared to elsewhere. Our results also indicate the potential for immigration from the west into the Chinese and Indochinese peninsular populations. This route was previously discounted but may present a means for new insecticide resistance traits to enter the Chinese and Indochinese peninsular populations. More generally, our results highlight the promise that population genomics holds for understanding the connectivity of pest insect populations characterised by high rates of gene flow.

## General

We thank Dr. Z. LV for providing some insect samples from Philippines.

## Funding

This work was supported by the National Key R&D Program of China (2017YFD0200602), the National Natural Science Foundation of China (31101430) and the Fundamental Research Funds for the Central Universities of China (2662017JC006).

## Author contributions

P.H & A.M.M analysed the data and designed the figures. J.P.H and Y.M.W wrote the manuscript, with revisions and input from A.M.M, and G.H.W. X.H.C prepared libraries for sequencing. C.X.X and Y.M.W collected the samples. Y.M.W designed the research.

## Competing interests

There are no competing interests to declare.

## Data and materials availability

Raw sequence data to be deposited on NCBI Short Read Archive (SRA), VCF’s and intermediate analysis files will be stored at UQ espace (https://espace.library.uq.edu.au/) and made publicly available on publication.

## Supplementary Material

Supplementary Table S1. Collection locations and sampling dates of the populations used in this study.

Supplementary Table S2. Mean sequencing coverage across *snps* for each individual.

Supplementary Table S3. Pairwise F_ST_’s calculated using Weir & Cockerham 1984 as implemented in OutFLANK below the diagonal, with 95% bounds above the diagonal calculated following the bootstrapping procedure implemented in hierfstat with 1,000 replicates.

Supplementary Figure 1. Plot of the assignment of the northern populations to the three putative sources, Philippines (top centre), “Indochinese peninsular” (bottom left) and Tengchong (bottom right), using DAPC.

