## Supplementary Figure 1 for "Migration dynamics of an important rice pest: the brown planthopper (*Nilaparvata lugens*) across Asia – insights from population genomics"

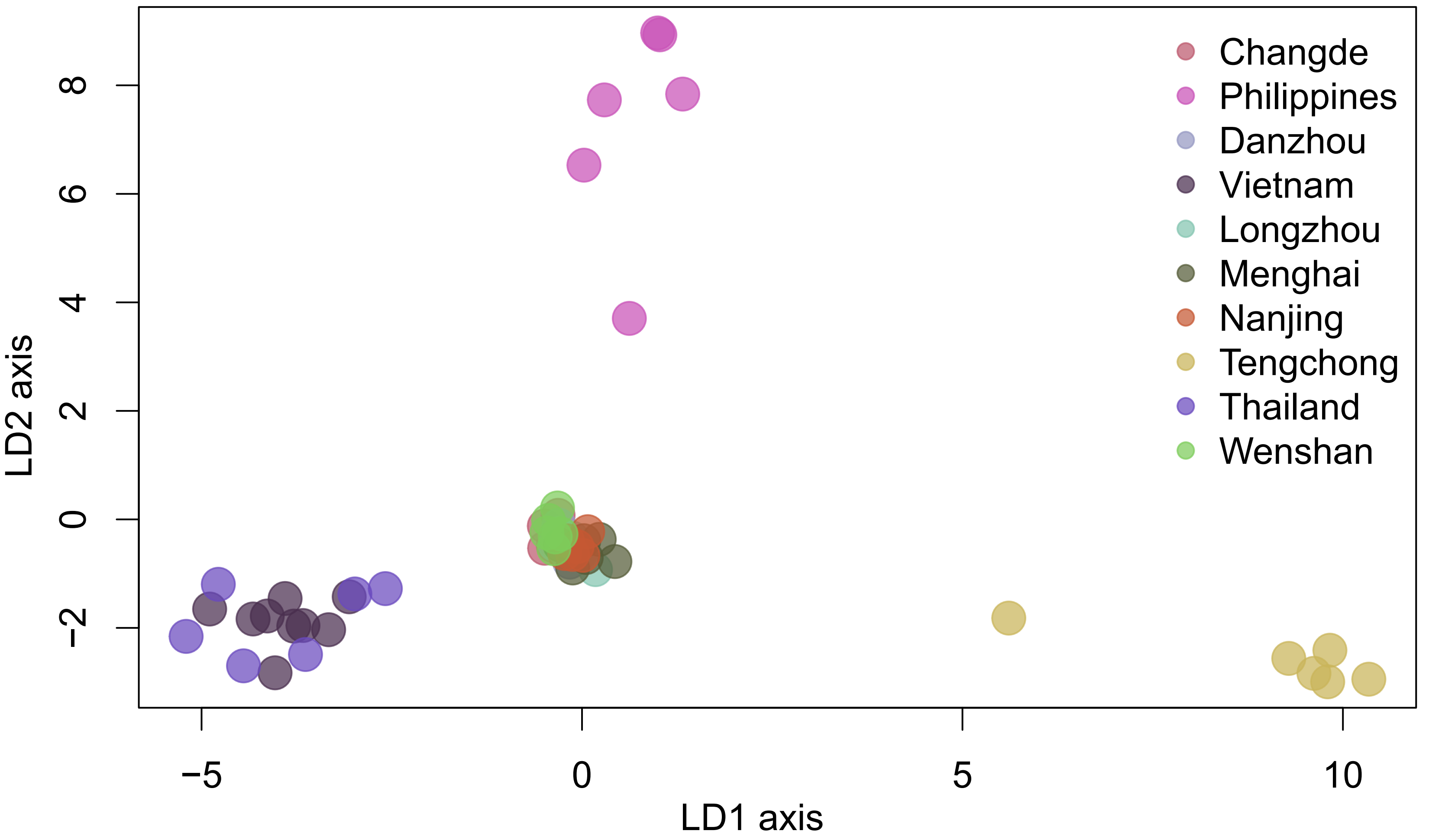


Supplementary Figure 1. Plot of the assignment of the northern populations to the three putative sources, Philippines (top centre), “Indochinese peninsular” (bottom left) and Tengchong (bottom right), using DAPC.
