## Supplementary Table 3 for "Migration dynamics of an important rice pest: the brown planthopper (*Nilaparvata lugens*) across Asia – insights from population genomics"

Supplementary Table 1. Pairwise $F_{ST}$'s calculated using Weir & Cockerham 1984 as implemented in hierfstat below the diagonal, with 95% bounds above the diagonal calculated following the bootstrapping procedure implemented in hierfstat with 1,000 replicates.

| | Changde | Philippines | Danzhou | Vietnam | Longzhou | Menghai | Nanjing | Tengchong | Thailand | Wenshan |
|---|---|---|---|---|---|---|---|---|---|---|
| Changde | | (0.0234 - 0.0242) | (0.001 - 0.0016) | (0.0072 - 0.0078) | (0.0021 - 0.0027) | (0.0039 - 0.0045) | (0.0067 - 0.0074) | (0.016 - 0.0167) | (0.0067 - 0.0074) | (0.0018 - 0.0024) |
| Philippines | 0.0238 | | (0.0196 - 0.0204) | (0.0206 - 0.0213) | (0.022 - 0.0228) | (0.0211 - 0.0219) | (0.0305 - 0.0314) | (0.0324 - 0.0333) | (0.021 - 0.0218) | (0.0189 - 0.0196) |
| Danzhou | 0.0013 | 0.0200 | | (0.0019 - 0.0025) | (0.0002 - 0.0008) | (0.0007 - 0.0012) | (0.0071 - 0.0077) | (0.0093 - 0.0099) | (0.0016 - 0.0021) | (0 - 0.0005) |
| Vietnam | 0.0075 | 0.0210 | 0.0022 | | (0.002 - 0.0025) | (0.002 - 0.0025) | (0.0101 - 0.0107) | (0.0121 - 0.0127) | (0.0018 - 0.0023) | (0.0045 - 0.005) |
| Longzhou | 0.0024 | 0.0224 | 0.0005 | 0.0023 | | (0.0014 - 0.002) | (0.005 - 0.0057) | (0.0094 - 0.01) | (0.003 - 0.0035) | (0.0021 - 0.0027) |
| Menghai | 0.0041 | 0.0215 | 0.0009 | 0.0022 | 0.0017 | | (0.0086 - 0.0093) | (0.0069 - 0.0075) | (0.0016 - 0.0021) | (0.0019 - 0.0024) |
| Nanjing | 0.0071 | 0.0309 | 0.0074 | 0.0104 | 0.0054 | 0.0089 | | (0.0182 - 0.0189) | (0.0117 - 0.0124) | (0.0083 - 0.009) |
| Tengchong | 0.0163 | 0.0329 | 0.0096 | 0.0124 | 0.0097 | 0.0072 | 0.0186 | | (0.0117 - 0.0124) | (0.0134 - 0.0141) |
| Thailand | 0.0070 | 0.0214 | 0.0018 | 0.0021 | 0.0032 | 0.0018 | 0.0121 | 0.0120 | | (0.0035 - 0.0041) |
| Wenshan | 0.0021 | 0.0193 | 0.0002 | 0.0047 | 0.0024 | 0.0022 | 0.0086 | 0.0137 | 0.0038 | |
